# Structural and Functional Disparities within the Human Gut Virome in terms of Genome Topology and Representative Genome Selection

**DOI:** 10.1101/2023.09.18.558355

**Authors:** Werner P Veldsman, Chao Yang, Zhenmiao Zhang, Yufen Huang, Lu Zhang

## Abstract

Circularity confers protection to viral genomes where linearity falls short, thereby fulfilling the form follows function aphorism. Yet, a shift away from morphology-based classification towards molecular and ecological classification of viruses is currently underway within the field of virology. Recent years have seen drastic changes in the ICTV operational definitions of viruses, particularly for those tailed phages that inhabit the human gut. After the abolition of the order Caudovirales, these tailed phages are best defined as members of the class Caudoviricetes, with taxonomic uncertainty prevailing at more granular taxa. In order to determine the epistemological value of genome topology in the context of the human gut virome, we designed a set of seven experiments to assay the impact of genome topology and representative viral selection on biological interpretation. Using Oxford Nanopore long reads for viral genome assembly coupled with Illumina short read polishing, we show that circular and linear virus genomes differ remarkably in terms of genome quality, GC-skew, tRNA gene frequency, structural variant frequency, cross-reference functional annotation (COG, KEGG, Pfam, and TIGRfam), state-of-the-art marker-based classification, and phage-host interaction. The disparity profile furthermore changes during dereplication. In particular, our phage-host interaction results demonstrate that proportional abundances are incomparable without due regard for genome topology and dereplication threshold, which necessitates the need for standardized reporting. As a best practice guideline, we recommend that comparative studies of the human gut virome should report the ratio of circular to linear viral genomes (ΔCL) along with the dereplication threshold so that structural and functional metrics can be placed into context when assessing biologically relevant metagenomic properties such as proportional abundance.

## 1. Introduction

Viruses evade classification by virtue of their miniscule size and vast diversity. Two years ago, the International Committee on Taxonomy of Viruses (ICTV) abolished the concept of a single type species, instead defining a species as a monophyletic group with multiple properties that distinguishes it from other monophyletic groups in the same genus [1]. In other words, viral species classification was inverted so that the monophyletic group henceforth defined the individual rather than a specimen defining the group. One year later, the ICTV abolished three major morphologically defined tailed phage virus families (Podoviridae, Siphoviridae, and Myoviridae) as well as their containing order Caudovirales [2]. This was not the first time that a taxonomic order meant for tailed phage membership was disbanded (see the discussion of phage classification and the 50-year-old redundant Urovirales order in [3]). The latest disbandment of the Caudovirales reflected a shifting consensus in the scientific community that viruses are better represented by molecular and ecological properties than by morphology. They also reflect the transient nature of viruses, and the impracticalities of merging the disciplines of morphology and virology in the quest for finding an appropriate representative of a virus.

Structure, however, remains a property of epistemological value since form follows function. The latter aphorism, that originated in the field of architecture, but subsequently spread to other scientific disciplines including biology, alludes to the pivotal role that structure serves in our understanding of reality. Another point to bear in mind is that viral anatomy is decidedly different from that of other organisms due to the minimalistic nature of the virus. Like macromolecules, such as lipids and carbohydrates, viral features that are commonly considered morphological (e.g. the capsid and tail), are measured on the nanoscale. If these quasi morphological features, that delineate the virus from the outside world, were to somehow be removed, all that would remain is the viral genome. The structure (or topology) of the viral genome would thus become a singular source of structural information. Despite phage genome sequences being much less conserved than phage structural proteins [4], sequence-based phage classification is often preferred over structure-based phage classification. Examples of the pervasiveness of sequence-based analysis is not only seen in viral classification, but also in the widespread use of multiple sequence alignment (MSA) tools in gene and protein ortholog detection.

The distinction between circularity and linearity of phage genomes has not received the attention it deserves since, as late as 1998, an eminent review of tailed phage literature stated that, “The genome of tailed phages is typically a single molecule of linear dsDNA” [3]. The importance of genome topology is in part recognized by the 1971 created Baltimore Classification System (BCS) [5] under the binary single or double stranded nucleic acid attribute. Nonetheless, the BCS system never has and still does not consider whether the viral genome is circular or linear. There are however ample examples in the literature of biochemical studies on polynucleotide strands that can serve as support for the argument that viruses should be classified by circular and linear topology. For example, circular and linear DNA have been shown to differ in their mechanism of cytoskeletal transport [6], in their anisotropy [7], and in their structural transition (as discussed in [8]). These proven biochemical distinctions between circular and linear DNA strongly suggest biologically relevant distinctions between circular and linear viral genomes. Taking the latter suggestion as a working hypothesis, we chose both extrinsic and intrinsic biologically relevant properties of viral DNA sequences as a basis for comparing circular and linear viral genomes. After considering typical measurements of interest in microbial analysis, we chose as extrinsic properties phage-host interaction, cross-reference functional annotation, and taxonomic classifiability, while we chose gene content, nucleotide frequency, dinucleotide skew, point and structural variation, and assembly quality as intrinsic properties. Moreover, our approach was multi-faceted in that we considered the preceding properties as dependent variables of molecular relatedness. In this manner, we could determine their values at intervals of average nucleotide identity to elucidate trends in biological interpretability during increasingly stringent rounds of representative virus selection.

The process of representative selection is in essence a clustering exercise in which viruses are grouped together based on a predefined genomic similarity criterium. At the strain level, no clustering is required. At the species level, 95% average nucleotide identity (ANI) is commonly considered appropriate. However, the genus and family ANI thresholds vary widely and can fall anywhere between 50% and 95%. With this non-standardized approach to delineating taxonomic boundaries at the genome-wide level for viruses, it is easy to appreciate the difficulties researchers encounter [9] when applying gene-level phylogenetic techniques that were honed on more evolutionarily stable genomes of living organisms. Moreover, these shortcomings in sequence analysis suggest that structural information should have a prominent role in virology. Considering that circularity versus linearity is not employed as a source of distinction by the major viral classification systems, and that there are no clear-cut ANI thresholds that define a virus taxon, a natural question that arises is: what impact does viral genome topology and dereplication thresholds have on structural and functional annotation? The results of our study that was aimed at addressing this question show that genomes classed by topology and dereplication stringency differ remarkably in terms of genome quality, GC-skew, tRNA gene frequency, structural variants, cross-reference functional annotation (COG, KEGG, Pfam, and TIGRfam), state-of-the-art marker-based classification, and phage-host interaction. Based on these findings, we recommend that best-practice comparative viromics of the human gut genome should always report the ratio of circular to linear viral genomes (ΔCL) along with the dereplication threshold so that molecular (e.g. gene frequency) and ecological (e.g. phage-host interaction) metrics can be accurately compared.

## 2. Methods

### 2.1 Source of human gut metagenomic sequencing reads

We relied on metagenomic sequencing datasets that were created during a previous study into genetic variation within the human gut microbiome [10]. These datasets were deposited at the NIH sequence read archive under BioProject PRJNA820119. We downloaded long read datasets for 200 Chinese individuals that were derived from an Oxford Nanopore Technology (ONT) PromethION platform via the EMBL-EBI FTP server (ftp.sra.ebi.ac.uk). We also downloaded 200 matching short read datasets (150 bp paired-end reads) derived from an Illumina Novaseq platform via the same EMBL-EBI server. Short reads were obtained with the purpose to polish assembled viral contigs. Summary long and short read statistics and plots were generated with NanoPlot v1.41.0 [11] and fastp v0.23.4 [12] to ascertain and compare the quality of the ONT long read and Illumina short read sequences.

### 2.2 Viral genome assembly, genome dereplication, and genome quality ascertainment

Viral genomes were assembled with viralFlye v0.2 [13], which requires as input contigs specifically generated by metaFlye [14]. We first passed raw ONT reads to the metaFlye v2.9.2-b1786 assembler using the nano-raw flag. To determine the assembly approach that would lead to the highest number of assembled viral genomes, we benchmarked the viralFlye assembler by permuting usage of (i) the metaFlye meta flag which is used to correct for uneven read coverage, (ii) internal short-read polishing using viralFlye, and (iii) external long read polishing using Medaka v1.6.1 (designed by ONT). To remove technical replicates that arise from repeated viral assembly for each of the 200 samples and to select representatives of the lower taxa, we carried out dereplication using the cluster functionality implemented in MMseqs2 v14.7e284 [15] with relaxed overlap calling (--cov-mode 1 -c 0.01). Another clustering program called dRep [16] is commonly used in microbial genome clustering, however, the authors of dRep state that virus genome clustering with their software requires usage of an independent genome completeness estimator. To promote genome completeness in our approach, we explicitly set the MMseqs2 cluster mode flag to 2 in order to reduce selection of shorter sequences as representative sequences. MMseqs2 automatically outputs representative sequences after clustering and we used all outputted sequences as representative sequences in downstream analysis. Minimum sequence identity (--min-seq-id) was set to 0.95 to dereplicate to species level, 0.70 to dereplicate to genus level, and 0.50 to dereplicate to family level. A threshold of 95% is commonly chosen as a species level cut-off (see [17] as an example) and is the Minimum Information about an Uncultivated Virus Genome (MIUVIG) standard for viral operational taxonomic units (vOTUs) [18]. Values between 50% and 95% are more arbitrarily selected in the literature as taxonomic boundaries. We chose a relatively stringent interpretation of 70% for genus and 50% for family since the assembled viral genomes were recalcitrant to clustering at higher sequence identities. Summary viral genome statistics (including GC-skew) were calculated using the fx2tab functionality in Seqkit v2.3.0 [19], and visualized using R 4.2.2 [20] with the library ggpubr v0.6.0 [21]. We assessed the quality of assembled viral constructs using checkV v1.0.1 [22]. The file containing the mean GC-skew and mean GC content that were calculated for each short read in all 200 samples was down sampled to 10% of its original size using simple random sampling in order to carry out comparative statistical procedures with the computational resources at our disposal. We noticed that the shape of the GC-skew distribution qualitatively differs between linear and circular sequences and we therefore used empirical cumulative distribution function analysis, bundled in the R package *twosamples* [23], to quantify the differences.

### 2.3 Viral genome annotation, taxonomic classification, and host prediction

The viral genomes, that were automatically classified as either circular or linear by the viralFlye assembler, were functionally annotated using geNomad v1.5.2 [24] with its end-to-end pipeline which includes a neural network implementation and custom viral profile database for the identification of proviruses and plasmids, for marker-based taxonomic classification, and for the functional annotation of viral genomes with cross-reference identifiers (COG, KEGG, Pfam, and TIGRfam). The most likely hosts for each assembled viral genome was predicted using iPHoP v1.3.2 [25]. Transfer RNAs (tRNAs) were detected using tRNAscan-SE v2.0.12 [26] with its general tRNA model selected as the tRNA detection model. Structural variants (including deletions (DELs), duplications (DUPs), inversions (INVs), insertions (INSs), and translocations) were detected using Sniffles v2.0.7 [27,28] with preprocessing using minimap2 v2.26-r1175 [29] and Samtools v1.17 [30]. We chose Sniffles because it detected a more diverse range of both real and simulated SVs (DELs, DUPs, INVs and INSs) compared to other long-read specific SV callers during a 2019 benchmark study [31]. Since read quality is especially important during variant calling, the long reads were filtered (q=12, u=5), trimmed (f=10; b=10,000) and deduplicated with fastp v0.23.4 [12] prior to mapping the reads to the viral genomes.

### 2.4 Protein structure prediction and ortholog detection

Determining protein orthology in terms of tertiary structure allows for the detection of remote homologs and analogs that are characterized by reduced sequence similarity. To supplement geNomad sequence based viral gene and protein prediction, we carried out structural orthology analysis using FoldSeek v 7.04e0ec8 [32], which is a newly developed tool capable of carrying out previously infeasible all-against-all comparisons of vast sets of tertiary protein structures. We first predicted the tertiary structures of the predicted viral protein sequences using API calls to the ESM Metagenomic Structure Atlas [33]. Resultant protein structure files in Protein Data Bank (PDB) format were then compared to tertiary structures in the Alphafold Protein Structure Database [34,35]. API calls to the ESM Metagenomic Structure Atlas did not robustly respond to requests. Structural orthologs were therefore not used in our comparison of topological and dereplication disparities, but solely to supplement sequence based ortholog detection.

## 3. Results and Discussion

### 3.1 Viral genome assembly quality and provirus detection

CheckV reported that the viral genomes that were assembled and polished with viralFlye are of good quality (Table 1). We followed a stringent approach in our quality assessment. We placed all viral genomes that were not deemed as high quality by both MIUVIG and CheckV standards into a low-quality category. Medium quality viral genome assemblies were therefore also placed in the low-quality category. The majority of viral genomes (96%) have no detectable host integration signals, of which 84% are of high quality. The latter percentage high-quality genomes remain consistent throughout dereplication. However, prior to dereplication, high quality circular genomes without detectable integration signals are 3.3x more abundant than high quality linear genomes. The ratio of high-quality circular to linear genomes, which we acronymize as ΔCL, decreases during dereplication to 2.1x at the species level, and further decreases to 1.6x at both the genus and family levels. The opposite trend is seen for high quality linear genomes that do have detectable host integration signals. For the latter presumed proviruses, high quality linear genomes outnumber circular genomes nearly fourfold, with the ratio increasing during dereplication.

**Table 1:** Quality of assembled viral genomes.

| Dereplication set | Topology | | Total | ΔCL | Quality | Integrated |
| --- | --- | --- | --- | --- | --- | --- |
|  | Circular | Linear |  |  |  |  |
| Strain | 938 | 281 | 1219 | 3.3 | high | No |
|  | 47 | 167 | 214 | 0.3 | low | No |
|  | 7 | 27 | 34 | 0.3 | high | Yes |
|  | 3 | 15 | 18 | 0.2 | low | Yes |
| Species | 352 | 167 | 519 | 2.1 | high | No |
|  | 22 | 79 | 101 | 0.3 | low | No |
|  | 3 | 24 | 27 | 0.1 | high | Yes |
|  | 1 | 9 | 10 | 0.1 | low | Yes |
| Genus | 165 | 105 | 270 | 1.6 | high | No |
|  | 12 | 34 | 46 | 0.4 | low | No |
|  | 2 | 16 | 18 | 0.1 | high | Yes |
|  | 0 | 5 | 5 | 0.0 | low | Yes |
| Family | 158 | 99 | 257 | 1.6 | high | No |
|  | 14 | 29 | 43 | 0.5 | low | No |
|  | 2 | 12 | 14 | 0.2 | high | Yes |
|  | 0 | 5 | 5 | 0.0 | low | Yes |

### 3.2 GC-skew is a biologically relevant property in topological genome conformation

The 200 ONT long read datasets from the Chen *et al* study contained a combined total of 147,299,829 reads with about a third of the reads having a quality score in excess of Q12 (Supplementary File S5). The long reads also had a mean GC content of 45.3% and a mean GC-skew of +0.13, indicating a slightly higher mean abundance of Guanine compared to Cytosine.

Fastp analysis of the Illumina short reads confirmed that read adapters had previous been trimmed and that 92.5% of the 11.2 billion reads (forward & reverse) across the 200 samples had a quality score in excess of Q30 (PHRED), likewise indicating previous quality filtering (Supplementary File S6). The short reads had a mean GC content of 46.8% and a mean GC-skew of +0.22, which as in the case of the long reads, indicates a slightly higher mean abundance of Guanine compared to Cytosine.

We determined that the best assembly approach to follow is viralFlye internal short-read polishing with the meta flag activated during the prerequisite metaFlye step (Table S1). The latter approach remains advantageous when considering the total number of genomes (circular and linear) retained after all three dereplication rounds, that is, at a minimum sequence identity of 0.95 (species level), 0.70 (genus level) and 0.50 (family level). However, after first and second round dereplication, more circular viral genomes are obtained when not using viralFlye internal short-read polishing. Nonetheless, in both these subcases our selected approach performed second best out of the five tested approaches, leaving our selected approach as the best approach in six out of the eight subcases (see *Most genomes in dereplication category* column in Table S1).

Statistical tests for normality of the mean GC content and mean GC-skew of both the long reads and short reads revealed that not one of the four respective vectors are normally distributed (Anderson-Darling, p<<0.05), and that GC content deviates at least ten times more from normality than GC-skew (as indicated by the Anderson-Darling test statistic). We accordingly tested for homogeneity of variance using a non-parametric test that is also robust against differences in sample size. Both the variance of GC content and the variance of GC-skew differed between long and short reads (Fligner-Killeen, p<<0.05), but in contrast to the greater departure from normality that was seen in GC content during the tests for normality, the greater departure from equal variance was between that of the GC-skew of long reads and the GC-skew of short reads (as indicated by the Fligner-Killeen test statistic). These tests served to empirical confirm that short reads have a higher frequency of Guanine than long reads, perhaps as a result of differing accuracies between long and short reads. Short reads are widely known to be more accurate than long reads, which leads to differences in genome assembly quality. With these statistics on metagenomic reads, we next analyzed GC-skew profiles in the assembled viral genomes.

In total, 1,485 viral genomes were assembled with the viralFlye assembler using our selected approach (Table 2) Roughly two-thirds of the viral constructs are circular and the rest are linear. As mentioned earlier, the ratio of circular to linear constructs decreases during dereplication, but circular constructs remain the most abundant. In contrast, the average length of viral genomes increases during dereplication, which suggests that manual adjustment of the MMseqs2 cluster-mode parameter promoted selection of longer representative viral genomes as expected.

**Table 2:** Summary statistics of the assembled viral genomes.

| Dereplication set | Number of genomes |  |  | Minimum length (bp) |  | Maximum length (bp) |  | Average length (bp) |  |
| --- | --- | --- | --- | --- | --- | --- | --- | --- | --- |
|  | Linear | Circular | Total | Linear | Circular | Linear | Circular | Linear | Circular |
| Strain | 490 | 995 | 1,485 | 5,446 | 5,301 | 232,946 | 213,711 | 58,213 | 67,205 |
| Species | 279 | 378 | 657 | 5,773 | 5,301 | 232,946 | 213,711 | 56,041 | 60,639 |
| Genus | 160 | 179 | 339 | 7,455 | 5,525 | 232,946 | 213,711 | 62,142 | 71,731 |
| Family | 145 | 174 | 319 | 7,455 | 5,525 | 232,946 | 213,711 | 63,799 | 72,772 |

The shape of the GC-skew density distribution of circular viral genomes differ noticeably from that of linear genomes (Figure 1). Empirical cumulative distribution function analysis revealed that the probability that circular and linear viral genome GC-skew values are from different distributions is significant at 50% identity (p=0.017), somewhat insignificant at 70% identity (p=0.111), insignificant at 95% identity (p=0.625), and only marginally significant prior to dereplication (p=0.052). The mean GC-Skew of the 1,485 genomes in the strain set was -0.09, while the dereplicated genomes had mean GC-skews of -0.48 (species set), -1.41 (genus set), and -0.71 (family set). Circular viral genomes consistently exhibit a higher mean Guanine abundance than linear genomes: strain set (circular: +0.23; linear: -0.73), species set (circular: -0.36; linear: -0.64), genus set (circular: -0.46; linear: -2.47), and family set (circular: +0.46, linear -2.12).

**Figure 1:**
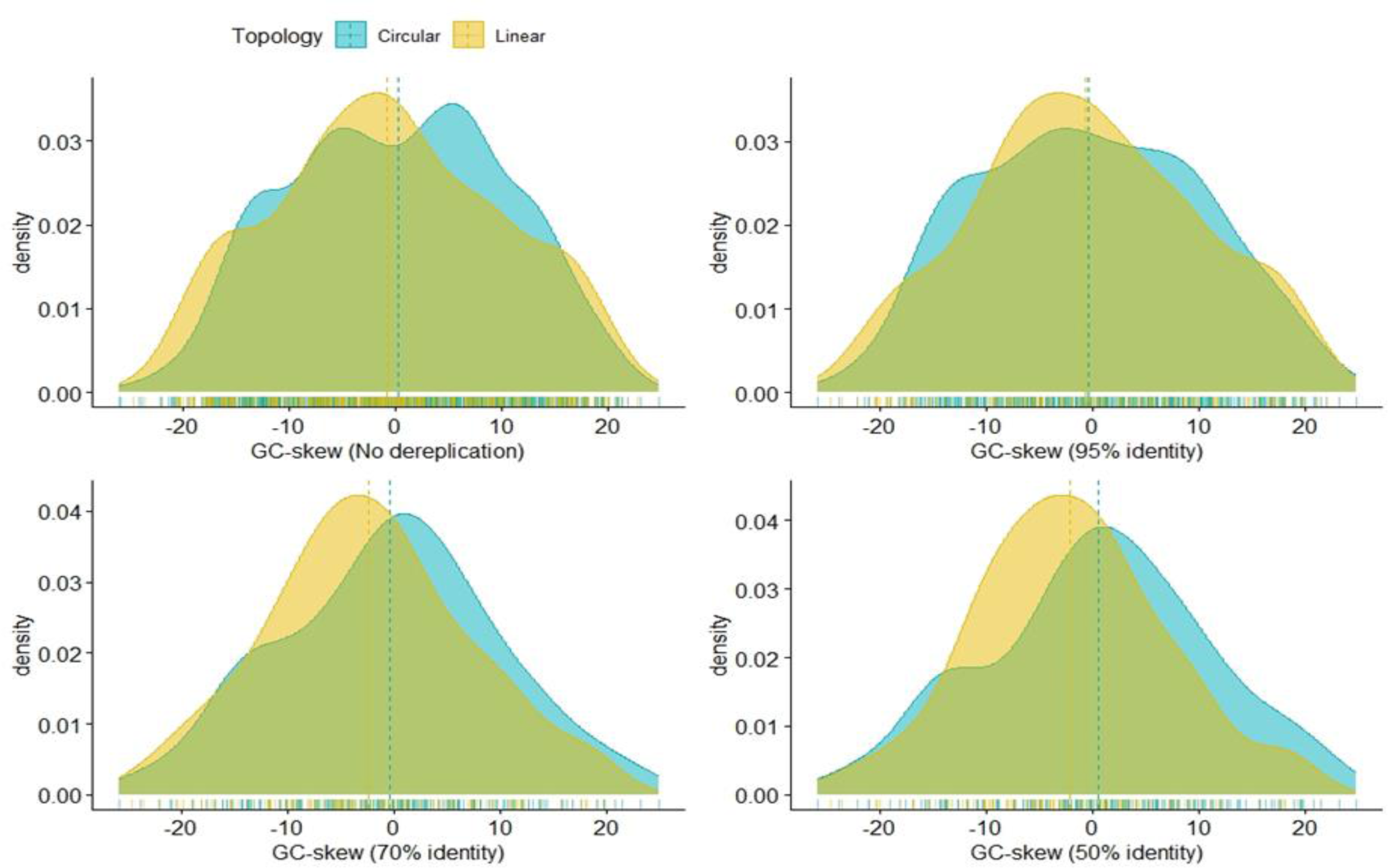
The importance of GC-skew in viral genome topology. (Top left to bottom right) Circular and linear genome GC-skew prior to dereplication and after 95%, 70% and 50% sequence similarity clustering. The shapes of the circular and linear GC-skew density distributions are noticeably different at each dereplication level. Empirical cumulative distribution function analysis (α=0.05) to determine whether circular and viral genome GC-skews come from the same distribution revealed that the two distributions are significantly different at 50% identity (p=0.017), somewhat insignificantly different at 70% identity (p=0.111), insignificantly different at 95% identity (p=0.625), and only marginally insignificantly different prior to dereplication (p=0.052).

GC-skew is known to be a non-trivial property that on occasion reflect the presence of certain genomic features. For example, a change of polarity (the sign) of GC-skew indicates such features as the origin of replication [36] and the site of mobile genetic element insertion [37]. The consistently higher mean GC-skew that we observe in circular viruses compared to linear viruses may have a role in structural configuration given that GC-skew in this case is a property that discriminates between two topological classes. Considering that these viral genomes were all assembled with long reads using short read polishing, it can be inferred that the difference in quality between long and short reads does not have bearing on the assembly of topologically different viral genomes exhibiting different GC-skews. In other words, the difference in GC-skew between circular and linear viral genomes cannot be an artifact caused by nucleotide base frequency and quality disparities between long and short reads. GC-skew is therefore a biologically relevant property in topological genome conformation.

### 3.3 Circular viral genomes contain more tRNAs than linear viral genomes

Phage genomes are known to contain tRNA genes in greater abundance than any other genes involved in translation [38]. It is furthermore known that virulent phages contain more tRNAs than temperate phages, which implies that tRNA function goes beyond protein synthesis to impact the viral life cycle. We accordingly analyzed the occurrence of tRNAs in our assembled viral genomes to determine whether there are differences in the number of detected tRNAs between circular and linear viral genomes, and whether tRNA detection frequency is affected by dereplication. Since more circular genomes were assembled than linear genomes in our study, we normalized the number of detected tRNAs. Despite the normalization step, we found that circular genomes have more detectable tRNAs than linear genomes on average (Figure 2). This bias towards circular genomes is constituted mostly by the number of tRNAs that were called with high confidence by tRNAscan-SE. The difference between circular and linear genomes in terms of the number of detected pseudo-tRNAs is trivial. A second observation we made is that the number of detected tRNAs reduce by about two-thirds during strain to species dereplication while the bias towards tRNAs in circular genomes more than doubles. The doubling of the aforementioned bias towards circular genomes reduces somewhat during second and third round dereplication, but remains nearly double the number observed prior to dereplication. This drastic initial increase in detected tRNA during strain to species dereplication is a pattern that we also observe during detection of structural variants (see section 3.4) and taxonomic classification (see section 3.6). We deduce from the similarity of these patterns that tRNAs may interact with circular genomes in a strain dependent manner since the patterns as mentioned in section 3.4 and section 3.6 are Crassvirales strain dependent. This deduction is supported by our observation that the ratio of tRNA-containing circular Crassvirales genomes to tRNA-containing circular genomes that are lost by clustering during strain to species dereplication, is more than double that of the ratio of tRNA-containing linear Crassvirales genomes to tRNA-containing linear genomes that are lost during strain to species dereplication. To end our assay of tRNAs, we compared the compositional abundance of the anti-codons on the detected tRNAs (Table 3). We found that Met-tRNA is always the most abundant tRNA regardless of topology and dereplication set, while Val-tRNA is always least common in linear viral genomes and His-tRNA (except in the strain set) is always least common in circular viral genomes.

**Figure 2:**
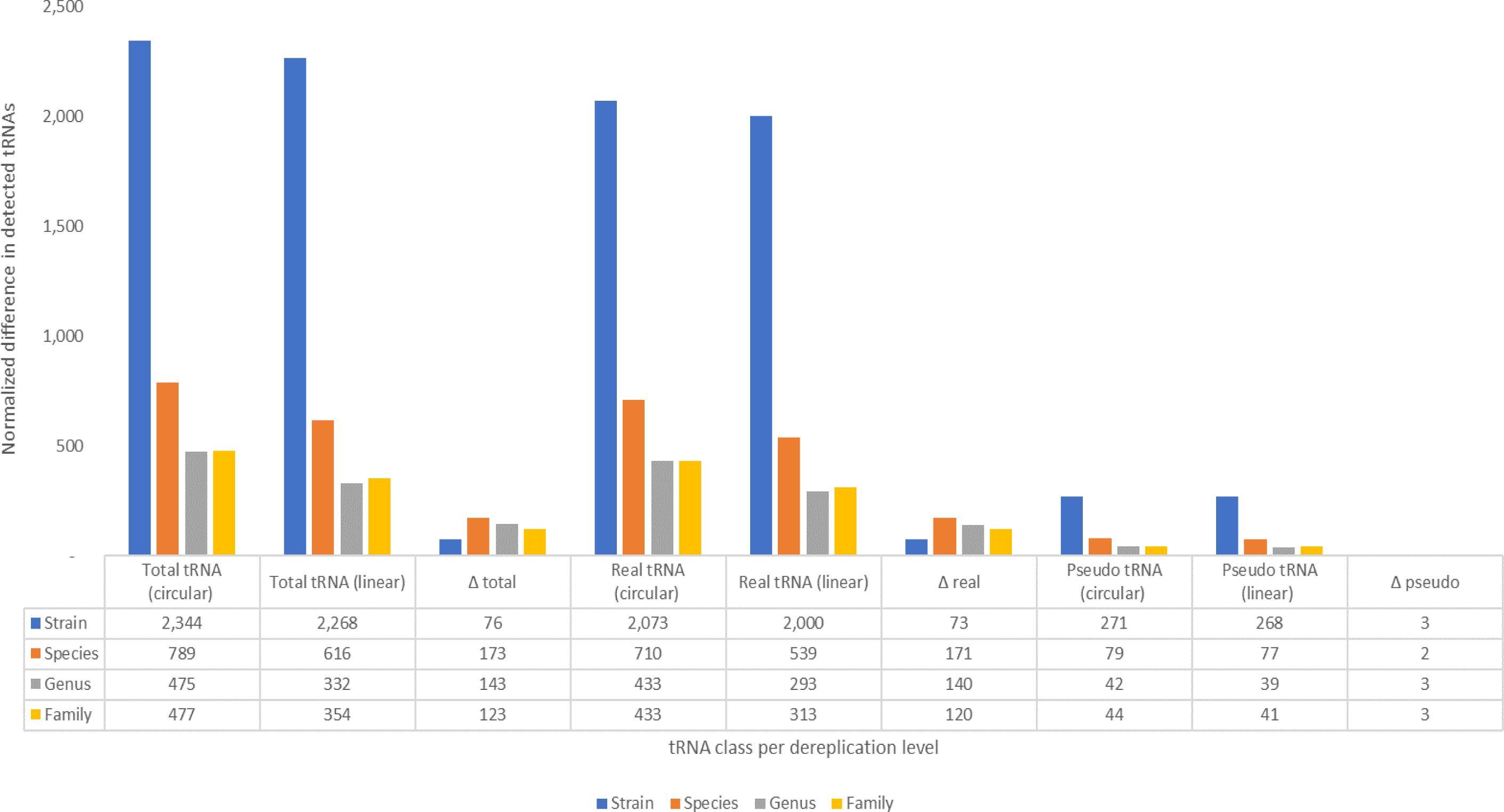
Transfer RNA frequency. Circular viral genomes contain a higher number of high confidences tRNAs than linear viral genomes, however, the difference (Δ) between circular and linear viral genomes in terms of pseudo tRNAs is marginal. The majority of detected tRNAs are lost during strain to species dereplication, while the bias towards circular viral genomes increases at the same time.

**Table 3:**
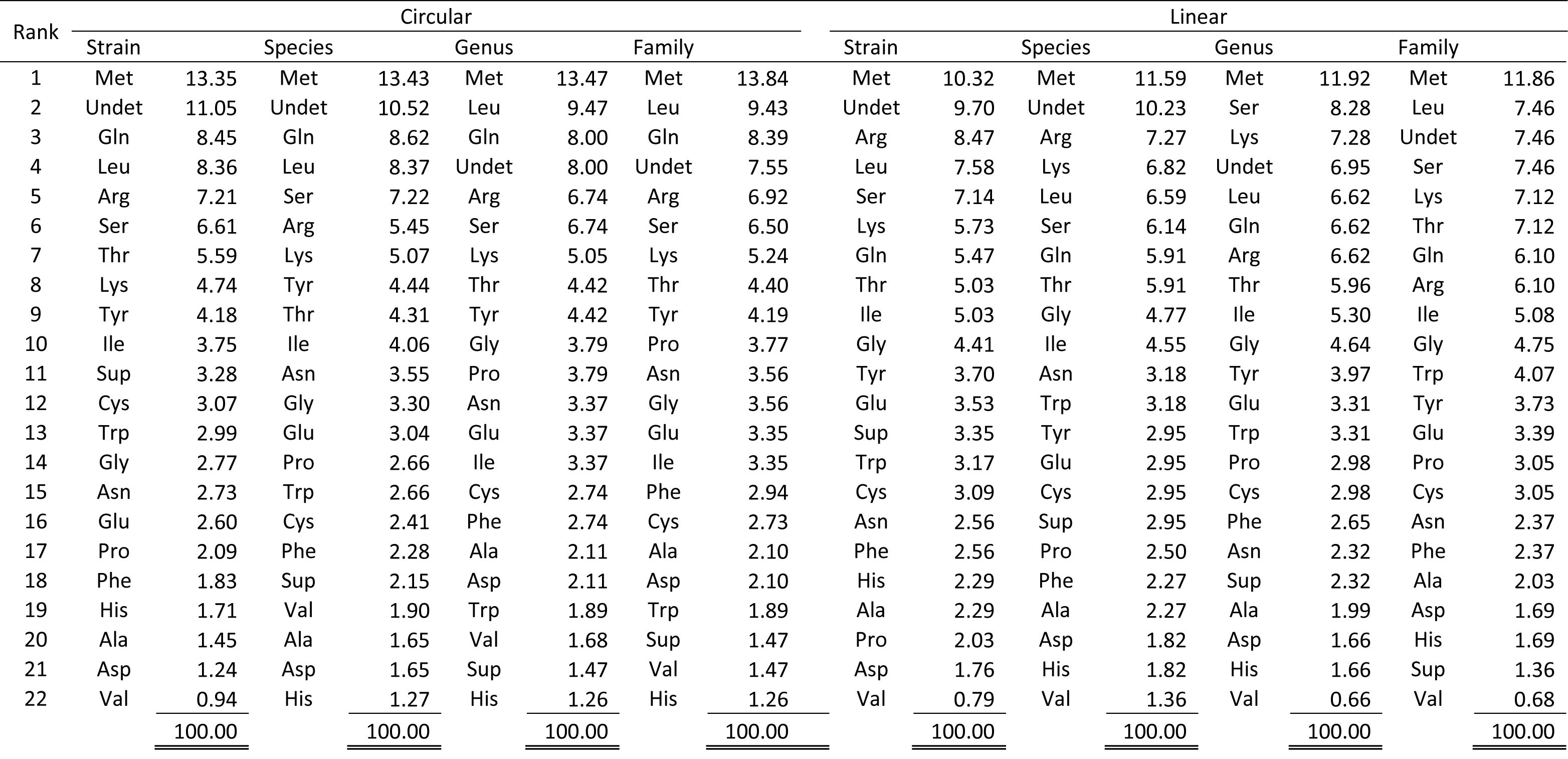
Ranked predicted tRNAs by their relative abundance (including pseudo tRNAs)

| Rank | Circular |  |  |  |  |  |  |  | Linear |  |  |  |  |  |  |  |
| --- | --- | --- | --- | --- | --- | --- | --- | --- | --- | --- | --- | --- | --- | --- | --- | --- |
|  | Strain |  | Species |  | Genus |  | Family |  | Strain |  | Species |  | Genus |  | Family |  |
| 1 | Met | 13.35 | Met | 13.43 | Met | 13.47 | Met | 13.84 | Met | 10.32 | Met | 11.59 | Met | 11.92 | Met | 11.86 |
| 2 | Undet | 11.05 | Undet | 10.52 | Leu | 9.47 | Leu | 9.43 | Undet | 9.70 | Undet | 10.23 | Ser | 8.28 | Leu | 7.46 |
| 3 | Gln | 8.45 | Gln | 8.62 | Gln | 8.00 | Gln | 8.39 | Arg | 8.47 | Arg | 7.27 | Lys | 7.28 | Undet | 7.46 |
| 4 | Leu | 8.36 | Leu | 8.37 | Undet | 8.00 | Undet | 7.55 | Leu | 7.58 | Lys | 6.82 | Undet | 6.95 | Ser | 7.46 |
| 5 | Arg | 7.21 | Ser | 7.22 | Arg | 6.74 | Arg | 6.92 | Ser | 7.14 | Leu | 6.59 | Leu | 6.62 | Lys | 7.12 |
| 6 | Ser | 6.61 | Arg | 5.45 | Ser | 6.74 | Ser | 6.50 | Lys | 5.73 | Ser | 6.14 | Gln | 6.62 | Thr | 7.12 |
| 7 | Thr | 5.59 | Lys | 5.07 | Lys | 5.05 | Lys | 5.24 | Gln | 5.47 | Gln | 5.91 | Arg | 6.62 | Gln | 6.10 |
| 8 | Lys | 4.74 | Tyr | 4.44 | Thr | 4.42 | Thr | 4.40 | Thr | 5.03 | Thr | 5.91 | Thr | 5.96 | Arg | 6.10 |
| 9 | Tyr | 4.18 | Thr | 4.31 | Tyr | 4.42 | Tyr | 4.19 | Ile | 5.03 | Gly | 4.77 | Ile | 5.30 | Ile | 5.08 |
| 10 | Ile | 3.75 | Ile | 4.06 | Gly | 3.79 | Pro | 3.77 | Gly | 4.41 | Ile | 4.55 | Gly | 4.64 | Gly | 4.75 |
| 11 | Sup | 3.28 | Asn | 3.55 | Pro | 3.79 | Asn | 3.56 | Tyr | 3.70 | Asn | 3.18 | Tyr | 3.97 | Trp | 4.07 |
| 12 | Cys | 3.07 | Gly | 3.30 | Asn | 3.37 | Gly | 3.56 | Glu | 3.53 | Trp | 3.18 | Glu | 3.31 | Tyr | 3.73 |
| 13 | Trp | 2.99 | Glu | 3.04 | Glu | 3.37 | Glu | 3.35 | Sup | 3.35 | Tyr | 2.95 | Trp | 3.31 | Glu | 3.39 |
| 14 | Gly | 2.77 | Pro | 2.66 | Ile | 3.37 | Ile | 3.35 | Trp | 3.17 | Glu | 2.95 | Pro | 2.98 | Pro | 3.05 |
| 15 | Asn | 2.73 | Trp | 2.66 | Cys | 2.74 | Phe | 2.94 | Cys | 3.09 | Cys | 2.95 | Cys | 2.98 | Cys | 3.05 |
| 16 | Glu | 2.60 | Cys | 2.41 | Phe | 2.74 | Cys | 2.73 | Asn | 2.56 | Sup | 2.95 | Phe | 2.65 | Asn | 2.37 |
| 17 | Pro | 2.09 | Phe | 2.28 | Ala | 2.11 | Ala | 2.10 | Phe | 2.56 | Pro | 2.50 | Asn | 2.32 | Phe | 2.37 |
| 18 | Phe | 1.83 | Sup | 2.15 | Asp | 2.11 | Asp | 2.10 | His | 2.29 | Phe | 2.27 | Sup | 2.32 | Ala | 2.03 |
| 19 | His | 1.71 | Val | 1.90 | Trp | 1.89 | Trp | 1.89 | Ala | 2.29 | Ala | 2.27 | Ala | 1.99 | Asp | 1.69 |
| 20 | Ala | 1.45 | Ala | 1.65 | Val | 1.68 | Sup | 1.47 | Pro | 2.03 | Asp | 1.82 | Asp | 1.66 | His | 1.69 |
| 21 | Asp | 1.24 | Asp | 1.65 | Sup | 1.47 | Val | 1.47 | Asp | 1.76 | His | 1.82 | His | 1.66 | Sup | 1.36 |
| 22 | Val | 0.94 | His | 1.27 | His | 1.26 | His | 1.26 | Val | 0.79 | Val | 1.36 | Val | 0.66 | Val | 0.68 |
|  |  | <u>100.00</u> |  | <u>100.00</u> |  | <u>100.00</u> |  | <u>100.00</u> |  | <u>100.00</u> |  | <u>100.00</u> |  | <u>100.00</u> |  | <u>100.00</u> |

### 3.4 Dereplication increases the detection rate of structural variants

Genes within virus genomes are tightly packed due to an intense natural constraint on viral genome size. Nevertheless, viral genomes are imperfect and undergo rapid mutation, which introduces not only the commonly known point mutations, but also structural variants (SVs). Previous research has shown that SVs have an impact on viral plaque size and viral dissemination in a strain dependent manner [39]. Since the identification of a strain has as prerequisite the identification of a species, and by implication must be accompanied by some form of representative genome selection (a process known as dereplication), we sought to determine SV profiles at the same dereplication levels that we compared elsewhere in the current study, that is, at the strain level, species level, genus level, and family level. In our analysis of tRNA profiles, we saw a pattern in which first round dereplication (dereplication from strain to species level) was accompanied by a sharp increase of tRNA detection in favor of circular genomes followed by a slight tapering off during subsequent dereplication. Here, in our analysis of SVs, a similar pattern emerged. We detected 30 structural variants in the strain set. Upon species level dereplication, the number of SVs more than doubled to 65. The implication in our opinion is that SVs are part of the reference genomes at strain level. Once representative species are selected by the process of dereplication, SVs are no longer identical to sequences in the species set and are flagged as variants. This argument is supported by our observation that of the 45 SVs that emerged during strain to species dereplication, 26 were in a genome classified under Crassvirales. This is noteworthy since less than 5% of the genomes in the species set were classified as Crassvirales genomes, indicating that structural variation in Crassvirales is strain dependent. We furthermore assayed the mutual inclusivity of SVs across the dereplication sets (Figure 3), where we noticed that all SVs that are present in every dereplication set are from linear genomes (Table S2).

**Figure 3:**
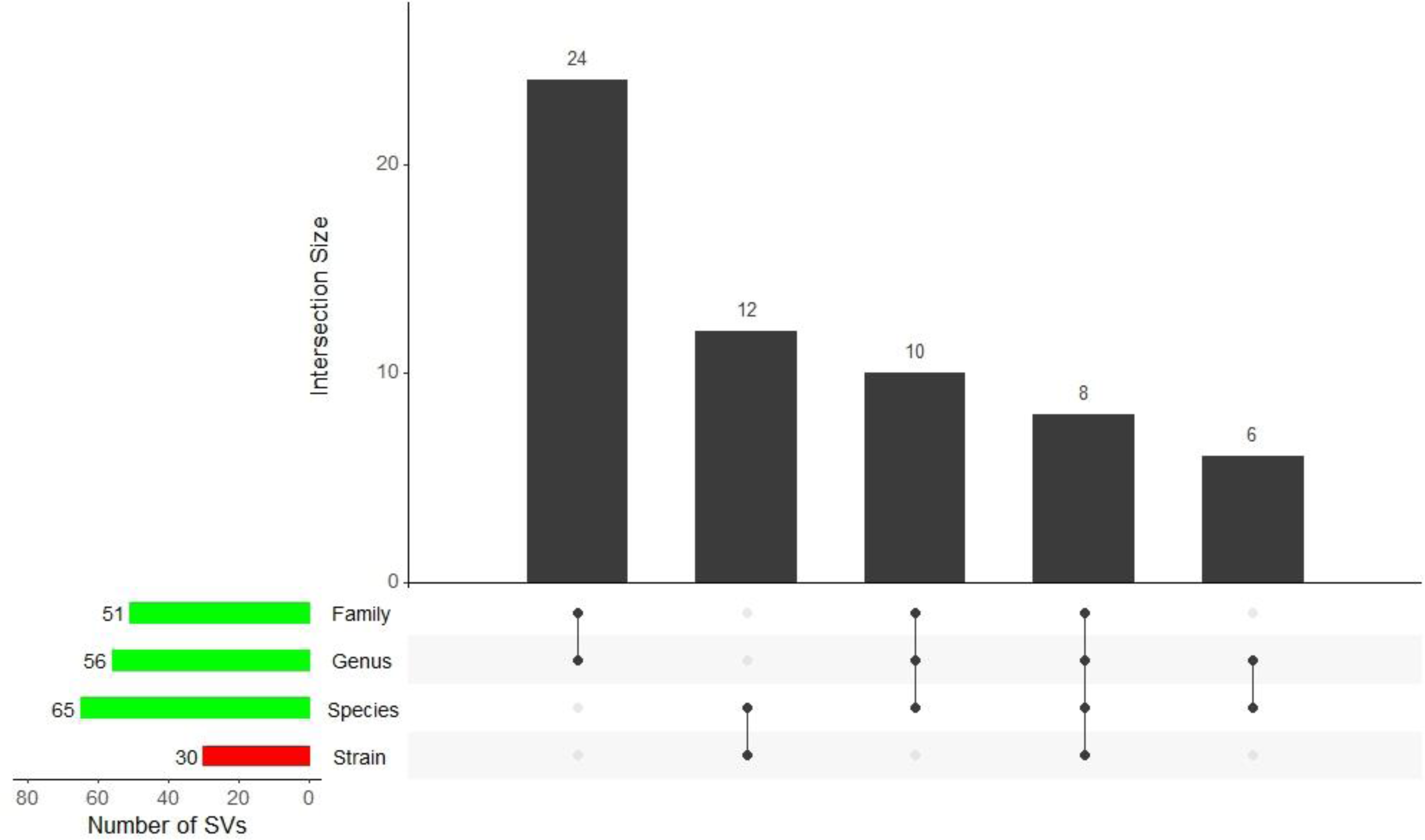
Structural variant frequency. Structural variants (SVs) are lower in the non-dereplicated strain set than in the dereplicated species set despite the latter containing less viral genomes. Only eight SVs appear consistently in all sets, with all eight detected in linear viral genomes.

### 3.5 Viral genome topology and representative genome selection affects functional annotation

Four databases were cross-referenced for functional annotations: Kyoto Encyclopedia of Genes and Genomes (KEGG) [40], Protein Families Database (Pfam) [41], The Institute of Genomic Research Functional Analysis and Classification of Proteins (TIGRfam) [42], and Clusters of Orthologous Groups (COG) [43]. Analysis of the annotations per topology and per dereplication level indicated that viral genome topology and dereplication strategy have a major impact on functional annotation (Table S3). We used the top 10 most frequent annotations as a metric to compare the relative frequency of annotations across different groups. Only three functional cross-references (“xrefs”) appear consistently in the top 10 annotations across topologies and across the dereplication levels: TIGR01547 (phage terminase), TIGR00673 (cyanase involved in cellular detoxification), and COG1783 (phage terminase). A fourth xref, PF05133 (phage portal protein), appears in all but one of the eight strata. TIGR01725 (phage morphogenesis) and COG5005 (Mu-like prophage protein) enter the top 10 xrefs during the first circular viral genome dereplication round, while two bacterial DNA primases (TIGR01391 and COG0358) drop to 53^rd^ and 78^th^ positions, respectively. The second round of dereplication promotes the relative frequency of annotation of PF13392 (HNH endonuclease) and PF03864 (phage major capsid protein E), while improving the position of the previously mentioned DNA primases (TIGR01391 and COG0358) to 43^rd^ and 27^th^, respectively. The final round of circular genome dereplication only had the effect of internally shuffling the top 10 xrefs, likely because the difference in number of genomes between the last two dereplication sets is not as great as the difference in the number of genomes between the first two dereplication round sets. A repeat of the analysis on linear genomes revealed that strain level linear genomes share only half of its top 10 xrefs with circular genomes. The first round of dereplication of linear genomes promoted the importance of TIGR01633 (putative phage tail component) and COG4926 (phage-related protein), while further dereplication of linear genomes had a less pronounced impact on the top 10 xrefs. An example of the implication of these observed differences in the relative frequency of xref annotation is that if a genome assembler is prone to assembling more circular genomes than linear genomes or vice versa, such technical properties of the assembler will propagate to functional analysis of the genomes where it will have a non-trivial impact on biological interpretation regardless of whether the assemblies are correct or not. Similarly, the process by which a representative sequence is selected will also have a non-trivial impact on downstream biological interpretation, which in turn has implications for viral strain analysis.

### 3.6 The vast majority of human gut viruses are tailed phages that defy marker-based classification

More than 99% of the viralFlye assemblies were confirmed as viral by geNomad taxonomic classification. The confirmed virus percentage decreased to >98% during first round dereplication and remained at that level for the remainder of the dereplication rounds. This decrease is not unexpected since non-viral representatives would necessarily be retained during dereplication. More than 98% of the viruses in the strain set belong to the realm Duplodnaviridia, which is defined as the double stranded DNA viruses that have in common a major capsid protein exhibiting a HK97 protein-fold. Here, too, dereplication decreased the Duplodnaviridia due to representatives of the Monodnaviria and Riboviria being detected. However, the share of Duplodnaviridia remained above 96% in all sets, and importantly, all Duplodnaviridia in all dereplication sets belonged to the class Caudoviricetes. The vast majority of Caudoviricetes were unclassifiable beyond the taxonomic rank of class (strain set = 88%; species set = 94%; genus set = 94%; family set = 93%). The reason why so many sequences are unclassifiable is due to geNomad’s stringent taxonomic classification approach in which at least 50% of a custom weighted score must support a specific taxon in order for a taxonomic name to be assigned to a genome. On the other hand, the 6% initial increase in unclassifiable genomes during strain to species dereplication is explained by classifiable CrAss-like phages going from having 165 strain representatives to having only 29 species representatives, thereby increasing the relative number of unclassifiable sequences. Although there are ample examples in the literature of attempts at phage family classification [44], there is still no standardized approach to evidence-based classification of metagenomic viruses [9]. Since the scope of our project is limited to determining the impact of genome topology and representative genome selection on taxonomic classification, we did not investigate the impact beyond the phylum and class taxonomic ranks, both of which are already clearly impacted by genome topology and dereplication. However, we suggest that a possible improvement on the limitations of geNomad’s MMseqs2-based protein-profile searches may lie in the use of tertiary protein structure comparison. As shown in the next section, tertiary structure prediction leads to the detection of plausible orthologs that challenge results derived from sequence-based ortholog detection.

### 3.7 The feasibility of phage-host comparative studies depends on the availability of strain data

Phages infect specific bacteria. The infection specificity is primarily determined by the specificity of adsorption, which correlates with specific receptors on the extracellular host surface (as discussed in [45]). However, once a phage attaches to a host, it must overcome a formidable molecular barrier to inject its DNA into the host cell. DNA topology confers physico-chemical properties that may play a role in this regard (as exemplified by the sought after properties of circular ssDNA in theranostics [46]). We therefore investigated whether there is a difference between the predicted bacterial hosts of linear and circular viral genomes. In line with our findings on structural and functional features, our results here revealed that genome topology also discriminates between linear and circular viral genomes in terms of their predicted bacterial hosts. In our strain set, circular viral genomes have a larger proportional difference between Fermicutes and Bacteroidota (referred to hereafter as ΔFB) than linear genomes (Figure 4). Dereplication analysis showed that the ΔFB for circular viral genomes decreases during dereplication with Fermicutes becoming less abundant and Bacteroidota becoming more abundant, while for linear genomes the ΔFB remains relatively stable albeit with lower abundance for both Firmicutes and Bacteroidata. The implication of the latter difference in ΔFB between circular and linear genomes is that robust comparisons cannot be made between metagenomic studies of the whole gut virome if there is not an explicit indication of the nucleotide similarity threshold that was used during representative genome selection. The lack of robustness is compounded by circular and linear viral genomes undergoing different changes in their ΔFBs during dereplication. We furthermore noticed an exception to the lowering of the proportion of Fermicutes during circular genome replication wherein the bacterial class Negativicutes (phylum: Fermicutes) was predicted more often as the host of phages with circular genomes than as the host of phages with linear genomes. The genus most frequently predicted as a host is Bacteroides sp., while the bacterial species most frequently associated with multiple high confidence Alphafold structural ortholog hits in the human gut virome is *Enterococcus faecium* (see data availability for protein models) – a bacterium whose clinical and non-clinal strains have distinct structural and functional features [47], suggesting a cryptic bacterial species that diverged in the absence of *in situ* ecological relationships.

**Figure 4:**
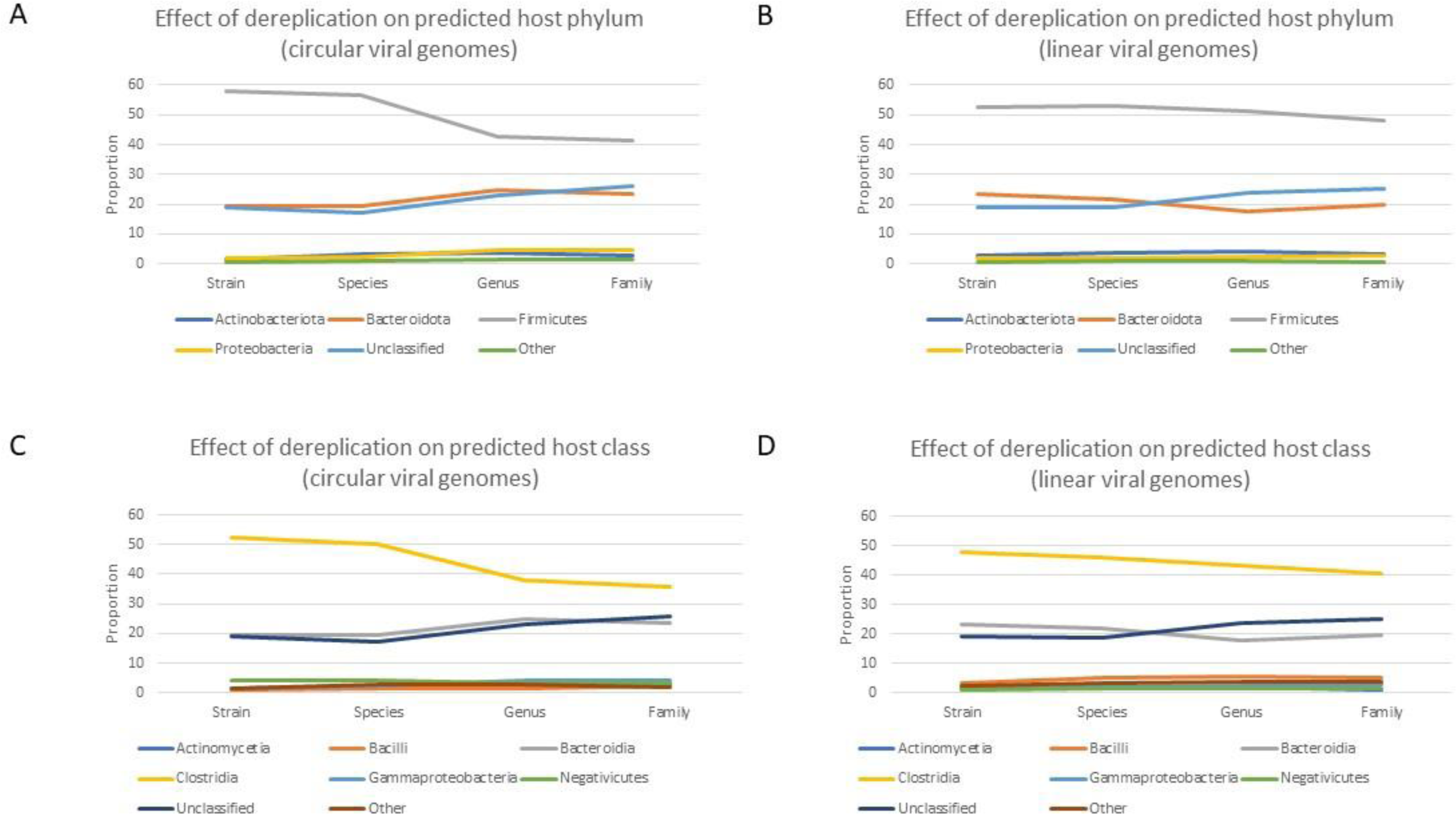
Predicted bacterial hosts. (A and C) Dereplication reduces the proportional difference between Firmicutes and Bacteroidota for circular viral genomes, while (B and D) the same proportional difference remains relatively stable during dereplication of linear viral genomes. These graphs furthermore clearly illustrate that dereplication threshold determines the proportional abundances of predicted bacterial host taxa in the human gut, which would have non-negatable impact on comparative studies of the virome.

## 4. Conclusion

Since viruses are nanobionts, drawing a line of distinction between which aspects of their study fall under morphology and which fall under molecular biological, is challenging. Although the concept of a type species is no longer recognized by the leading authority in virology, the concept of a monophyletic group that has replaced the type species, still requires shared molecular and ecological characteristics. In this study, we demonstrate that genome topology and representative genome selection have a non-trivial impact on biological interpretation. We report on the results of seven separate experiments to assay the difference between circular and linear genomes. Each experiment revealed that there is a remarkable difference between circular and linear viral genomes in terms of not only molecular features, but also in terms of the process by which representative viruses are selected. To allow for the accurate comparison of human gut viromes between studies, we recommend that researchers report the ratio of circular to linear viruses (ΔCL) along with dereplication thresholds. The ΔCL ratio is limited in that it does not model exceptions to observed differences between circular and linear viral genomes, such as that circular phages exhibit an overall decrease in the proportion of predicted Firmicute hosts during dereplication with the class Negativicutes being a notable exception. Nevertheless, the ΔCL ratio provides a basis for due consideration of the structural and functional differences between circular and linear viral genomes, and serves as draft for future modelling of the proportional abundance of circular and linear viruses in the human gut.

## Supporting information

All supplementary files

## List of Tables and Figures

Table 1: Quality of assembled viral genomes

Table 2: Summary statistics of the assembled viral genomes

Table 3: Ranked predicted tRNAs by their relative abundance (including pseudo tRNAs)

Figure 1: The importance of GC-skew in viral genome topology

Figure 2: Transfer RNA frequency

Figure 3: Structural variant frequency

Figure 4: Predicted bacterial hosts

## List of Supplementary Material

Table S1: The effect of polishing and the meta flag

Table S2: Pervasive structural variants

Table S3: Cross-reference functional annotations per topology per dereplication level

File S1: Strain set circular and linear viral genomes

File S2: Species set circular and linear viral genomes

File S3: Genus set circular and linear viral genomes

File S4: Family set circular and linear viral genomes

File S5: Long read summary statistics

File S6: Short read summary statistics

## Data availability

Interactive 3D protein models of the *Enterococcus faecium* structural orthologs can be viewed at https://alphafold.com under model IDs A0A132P7M2, A0A132Z4D1, A0A132Z369 and A0A133CLV7. Circular and linear virus genomes are included in supplementary File S1 through File S4 for the strain, species, genus and family sets. Circular and linear constructs in the preceding files are discernible by their FASTA header names. Scripting in this project was carried out using the R programming language. Code snippets are available at https://github.com/Werner0/tailed_phages. Long reads and short reads from the Chen et al. [10] study are hosted by the EMBL-EBI (see methods section).

## Author Contributions

All authors listed have made an intellectual contribution to the work and approved it for publication.

