## Supplementary material for "Structural and Functional Disparities within the Human Gut Virome in terms of Genome Topology and Representative Genome Selection": All supplementary files: File_S5_long_read_statistics.html

NanoPlot Report


### NanoPlot statistics report

#### Menu

- Summary Statistics
- Plots
  - Weighted histogram of read lengths
  - Weighted histogram of read lengths after log transformation
  - Non weighted histogram of read lengths
  - Non weighted histogram of read lengths after log transformation
  - Yield by length
  - Read lengths vs Average read quality plot using dots
  - Read lengths vs Average read quality kde plot
  - Read lengths vs Average read quality plot using dots after log transformation of read lengths
  - Read lengths vs Average read quality kde plot
- Report issue on Github


#### NanoPlot reports

##### Summary statistics

|  |  |
| --- | --- |
| General summary |  |
| Mean read length | 7,281.7 |
| Mean read quality | 10.1 |
| Median read length | 6,849.0 |
| Median read quality | 11.0 |
| Number of reads | 147,299,829.0 |
| Read length N50 | 8,774.0 |
| STDEV read length | 3,943.0 |
| Total bases | 1,072,596,978,744.0 |
| Number, percentage and megabases of reads above quality cutoffs |  |
| >Q5 | 147298188 (100.0%) 1072583.9Mb |
| >Q7 | 147099803 (99.9%) 1070908.7Mb |
| >Q10 | 100744081 (68.4%) 722355.2Mb |
| >Q12 | 45684254 (31.0%) 307676.8Mb |
| >Q15 | 7220993 (4.9%) 42072.6Mb |
| Top 5 highest mean basecall quality scores and their read lengths |  |
| 1 | 23.5 (2084) |
| 2 | 22.5 (5989) |
| 3 | 22.4 (5198) |
| 4 | 22.4 (2413) |
| 5 | 22.1 (4986) |
| Top 5 longest reads and their mean basecall quality score |  |
| 1 | 630912 (7.7) |
| 2 | 440030 (7.7) |
| 3 | 393688 (8.0) |
| 4 | 303639 (7.9) |
| 5 | 267901 (9.1) |

##### Plots

Weighted histogram of read lengths

###### Weighted histogram of read lengths

Weighted histogram of read lengths after log transformation

###### Weighted histogram of read lengths after log transformation

Non weighted histogram of read lengths

###### Non weighted histogram of read lengths

Non weighted histogram of read lengths after log transformation

###### Non weighted histogram of read lengths after log transformation

Yield by length

###### Yield by length

Read lengths vs Average read quality plot using dots

###### Read lengths vs Average read quality plot using dots

Read lengths vs Average read quality kde plot

###### Read lengths vs Average read quality kde plot

Read lengths vs Average read quality plot using dots after log transformation of read lengths

###### Read lengths vs Average read quality plot using dots after log transformation of read lengths

Read lengths vs Average read quality kde plot

###### Read lengths vs Average read quality kde plot
