## Supplementary material for "Structural and Functional Disparities within the Human Gut Virome in terms of Genome Topology and Representative Genome Selection": All supplementary files: File_S6_short_read_statistics.html

fastp report at 2023-08-01 17:46:56 

### fastp report Summary General | | | | --- | --- | | fastp version: | 0.23.4 (https://github.com/OpenGene/fastp) | | sequencing: | single end (150 cycles) | | mean length before filtering: | 150bp | | mean length after filtering: | 149bp | | duplication rate: | 74.102206% (may be overestimated since this is SE data) | | Detected read1 adapter: | AGATCGGAAGAGCACACGTCTGAACTCCAGTCA | Before filtering | | | | --- | --- | | total reads: | 11.275840 G | | total bases: | 1.691376 T | | Q20 bases: | 1.647705 T (97.418011%) | | Q30 bases: | 1.565156 T (92.537455%) | | GC content: | 46.842690% | After filtering | | | | --- | --- | | total reads: | 11.251165 G | | total bases: | 1.684814 T | | Q20 bases: | 1.643240 T (97.532448%) | | Q30 bases: | 1.561418 T (92.676023%) | | GC content: | 46.837537% | Filtering result | | | | --- | --- | | reads passed filters: | 11.251165 G (99.781170%) | | reads with low quality: | 24.046116 M (0.213253%) | | reads with too many N: | 8.811000 K (0.000078%) | | reads too short: | 619.986000 K (0.005498%) | Adapters Adapter or bad ligation of read1 The input has little adapter percentage (~0.171001%), probably it's trimmed before. | | | | --- | --- | | Sequence | Occurrences | | AGATC | 16326054 | | AGATCG | 8226061 | | AGATCGG | 6152932 | | AGATCGGA | 5528550 | | AGATCGGAA | 5243812 | | AGATCGGAAG | 5004275 | | AGATCGGAAGA | 4835126 | | AGATCGGAAGAG | 4689786 | | AGATCGGAAGAGC | 4490077 | | AGATCGGAAGAGCA | 2176126 | | AGATCGGAAGAGCG | 2177669 | | AGATCGGAAGAGCAC | 2107566 | | AGATCGGAAGAGCACA | 2026089 | | AGATCGGAAGAGCACAC | 1960795 | | AGATCGGAAGAGCACACG | 1899719 | | AGATCGGAAGAGCACACGT | 1813693 | | AGATCGGAAGAGCACACGTC | 1758803 | | AGATCGGAAGAGCACACGTCT | 1702610 | | AGATCGGAAGAGCACACGTCTG | 1629171 | | AGATCGGAAGAGCACACGTCTGA | 1576624 | | AGATCGGAAGAGCACACGTCTGAA | 1531217 | | AGATCGGAAGAGCACACGTCTGAAC | 1463910 | | AGATCGGAAGAGCACACGTCTGAACT | 1413445 | | AGATCGGAAGAGCACACGTCTGAACTC | 1372085 | | AGATCGGAAGAGCACACGTCTGAACTCC | 1317316 | | AGATCGGAAGAGCACACGTCTGAACTCCAGTC | 1653841 | | other adapter sequences | 41184818 | Before filtering Before filtering: read1: quality Value of each position will be shown on mouse over. Before filtering: read1: base contents Value of each position will be shown on mouse over. Before filtering: read1: KMER counting Darker background means larger counts. The count will be shown on mouse over. | | | | | | | | | | | | | | | | | | | --- | --- | --- | --- | --- | --- | --- | --- | --- | --- | --- | --- | --- | --- | --- | --- | --- | | | AA | AT | AC | AG | TA | TT | TC | TG | CA | CT | CC | CG | GA | GT | GC | GG | | AAA | AAAAA | AAAAT | AAAAC | AAAAG | AAATA | AAATT | AAATC | AAATG | AAACA | AAACT | AAACC | AAACG | AAAGA | AAAGT | AAAGC | AAAGG | | AAT | AATAA | AATAT | AATAC | AATAG | AATTA | AATTT | AATTC | AATTG | AATCA | AATCT | AATCC | AATCG | AATGA | AATGT | AATGC | AATGG | | AAC | AACAA | AACAT | AACAC | AACAG | AACTA | AACTT | AACTC | AACTG | AACCA | AACCT | AACCC | AACCG | AACGA | AACGT | AACGC | AACGG | | AAG | AAGAA | AAGAT | AAGAC | AAGAG | AAGTA | AAGTT | AAGTC | AAGTG | AAGCA | AAGCT | AAGCC | AAGCG | AAGGA | AAGGT | AAGGC | AAGGG | | ATA | ATAAA | ATAAT | ATAAC | ATAAG | ATATA | ATATT | ATATC | ATATG | ATACA | ATACT | ATACC | ATACG | ATAGA | ATAGT | ATAGC | ATAGG | | ATT | ATTAA | ATTAT | ATTAC | ATTAG | ATTTA | ATTTT | ATTTC | ATTTG | ATTCA | ATTCT | ATTCC | ATTCG | ATTGA | ATTGT | ATTGC | ATTGG | | ATC | ATCAA | ATCAT | ATCAC | ATCAG | ATCTA | ATCTT | ATCTC | ATCTG | ATCCA | ATCCT | ATCCC | ATCCG | ATCGA | ATCGT | ATCGC | ATCGG | | ATG | ATGAA | ATGAT | ATGAC | ATGAG | ATGTA | ATGTT | ATGTC | ATGTG | ATGCA | ATGCT | ATGCC | ATGCG | ATGGA | ATGGT | ATGGC | ATGGG | | ACA | ACAAA | ACAAT | ACAAC | ACAAG | ACATA | ACATT | ACATC | ACATG | ACACA | ACACT | ACACC | ACACG | ACAGA | ACAGT | ACAGC | ACAGG | | ACT | ACTAA | ACTAT | ACTAC | ACTAG | ACTTA | ACTTT | ACTTC | ACTTG | ACTCA | ACTCT | ACTCC | ACTCG | ACTGA | ACTGT | ACTGC | ACTGG | | ACC | ACCAA | ACCAT | ACCAC | ACCAG | ACCTA | ACCTT | ACCTC | ACCTG | ACCCA | ACCCT | ACCCC | ACCCG | ACCGA | ACCGT | ACCGC | ACCGG | | ACG | ACGAA | ACGAT | ACGAC | ACGAG | ACGTA | ACGTT | ACGTC | ACGTG | ACGCA | ACGCT | ACGCC | ACGCG | ACGGA | ACGGT | ACGGC | ACGGG | | AGA | AGAAA | AGAAT | AGAAC | AGAAG | AGATA | AGATT | AGATC | AGATG | AGACA | AGACT | AGACC | AGACG | AGAGA | AGAGT | AGAGC | AGAGG | | AGT | AGTAA | AGTAT | AGTAC | AGTAG | AGTTA | AGTTT | AGTTC | AGTTG | AGTCA | AGTCT | AGTCC | AGTCG | AGTGA | AGTGT | AGTGC | AGTGG | | AGC | AGCAA | AGCAT | AGCAC | AGCAG | AGCTA | AGCTT | AGCTC | AGCTG | AGCCA | AGCCT | AGCCC | AGCCG | AGCGA | AGCGT | AGCGC | AGCGG | | AGG | AGGAA | AGGAT | AGGAC | AGGAG | AGGTA | AGGTT | AGGTC | AGGTG | AGGCA | AGGCT | AGGCC | AGGCG | AGGGA | AGGGT | AGGGC | AGGGG | | TAA | TAAAA | TAAAT | TAAAC | TAAAG | TAATA | TAATT | TAATC | TAATG | TAACA | TAACT | TAACC | TAACG | TAAGA | TAAGT | TAAGC | TAAGG | | TAT | TATAA | TATAT | TATAC | TATAG | TATTA | TATTT | TATTC | TATTG | TATCA | TATCT | TATCC | TATCG | TATGA | TATGT | TATGC | TATGG | | TAC | TACAA | TACAT | TACAC | TACAG | TACTA | TACTT | TACTC | TACTG | TACCA | TACCT | TACCC | TACCG | TACGA | TACGT | TACGC | TACGG | | TAG | TAGAA | TAGAT | TAGAC | TAGAG | TAGTA | TAGTT | TAGTC | TAGTG | TAGCA | TAGCT | TAGCC | TAGCG | TAGGA | TAGGT | TAGGC | TAGGG | | TTA | TTAAA | TTAAT | TTAAC | TTAAG | TTATA | TTATT | TTATC | TTATG | TTACA | TTACT | TTACC | TTACG | TTAGA | TTAGT | TTAGC | TTAGG | | TTT | TTTAA | TTTAT | TTTAC | TTTAG | TTTTA | TTTTT | TTTTC | TTTTG | TTTCA | TTTCT | TTTCC | TTTCG | TTTGA | TTTGT | TTTGC | TTTGG | | TTC | TTCAA | TTCAT | TTCAC | TTCAG | TTCTA | TTCTT | TTCTC | TTCTG | TTCCA | TTCCT | TTCCC | TTCCG | TTCGA | TTCGT | TTCGC | TTCGG | | TTG | TTGAA | TTGAT | TTGAC | TTGAG | TTGTA | TTGTT | TTGTC | TTGTG | TTGCA | TTGCT | TTGCC | TTGCG | TTGGA | TTGGT | TTGGC | TTGGG | | TCA | TCAAA | TCAAT | TCAAC | TCAAG | TCATA | TCATT | TCATC | TCATG | TCACA | TCACT | TCACC | TCACG | TCAGA | TCAGT | TCAGC | TCAGG | | TCT | TCTAA | TCTAT | TCTAC | TCTAG | TCTTA | TCTTT | TCTTC | TCTTG | TCTCA | TCTCT | TCTCC | TCTCG | TCTGA | TCTGT | TCTGC | TCTGG | | TCC | TCCAA | TCCAT | TCCAC | TCCAG | TCCTA | TCCTT | TCCTC | TCCTG | TCCCA | TCCCT | TCCCC | TCCCG | TCCGA | TCCGT | TCCGC | TCCGG | | TCG | TCGAA | TCGAT | TCGAC | TCGAG | TCGTA | TCGTT | TCGTC | TCGTG | TCGCA | TCGCT | TCGCC | TCGCG | TCGGA | TCGGT | TCGGC | TCGGG | | TGA | TGAAA | TGAAT | TGAAC | TGAAG | TGATA | TGATT | TGATC | TGATG | TGACA | TGACT | TGACC | TGACG | TGAGA | TGAGT | TGAGC | TGAGG | | TGT | TGTAA | TGTAT | TGTAC | TGTAG | TGTTA | TGTTT | TGTTC | TGTTG | TGTCA | TGTCT | TGTCC | TGTCG | TGTGA | TGTGT | TGTGC | TGTGG | | TGC | TGCAA | TGCAT | TGCAC | TGCAG | TGCTA | TGCTT | TGCTC | TGCTG | TGCCA | TGCCT | TGCCC | TGCCG | TGCGA | TGCGT | TGCGC | TGCGG | | TGG | TGGAA | TGGAT | TGGAC | TGGAG | TGGTA | TGGTT | TGGTC | TGGTG | TGGCA | TGGCT | TGGCC | TGGCG | TGGGA | TGGGT | TGGGC | TGGGG | | CAA | CAAAA | CAAAT | CAAAC | CAAAG | CAATA | CAATT | CAATC | CAATG | CAACA | CAACT | CAACC | CAACG | CAAGA | CAAGT | CAAGC | CAAGG | | CAT | CATAA | CATAT | CATAC | CATAG | CATTA | CATTT | CATTC | CATTG | CATCA | CATCT | CATCC | CATCG | CATGA | CATGT | CATGC | CATGG | | CAC | CACAA | CACAT | CACAC | CACAG | CACTA | CACTT | CACTC | CACTG | CACCA | CACCT | CACCC | CACCG | CACGA | CACGT | CACGC | CACGG | | CAG | CAGAA | CAGAT | CAGAC | CAGAG | CAGTA | CAGTT | CAGTC | CAGTG | CAGCA | CAGCT | CAGCC | CAGCG | CAGGA | CAGGT | CAGGC | CAGGG | | CTA | CTAAA | CTAAT | CTAAC | CTAAG | CTATA | CTATT | CTATC | CTATG | CTACA | CTACT | CTACC | CTACG | CTAGA | CTAGT | CTAGC | CTAGG | | CTT | CTTAA | CTTAT | CTTAC | CTTAG | CTTTA | CTTTT | CTTTC | CTTTG | CTTCA | CTTCT | CTTCC | CTTCG | CTTGA | CTTGT | CTTGC | CTTGG | | CTC | CTCAA | CTCAT | CTCAC | CTCAG | CTCTA | CTCTT | CTCTC | CTCTG | CTCCA | CTCCT | CTCCC | CTCCG | CTCGA | CTCGT | CTCGC | CTCGG | | CTG | CTGAA | CTGAT | CTGAC | CTGAG | CTGTA | CTGTT | CTGTC | CTGTG | CTGCA | CTGCT | CTGCC | CTGCG | CTGGA | CTGGT | CTGGC | CTGGG | | CCA | CCAAA | CCAAT | CCAAC | CCAAG | CCATA | CCATT | CCATC | CCATG | CCACA | CCACT | CCACC | CCACG | CCAGA | CCAGT | CCAGC | CCAGG | | CCT | CCTAA | CCTAT | CCTAC | CCTAG | CCTTA | CCTTT | CCTTC | CCTTG | CCTCA | CCTCT | CCTCC | CCTCG | CCTGA | CCTGT | CCTGC | CCTGG | | CCC | CCCAA | CCCAT | CCCAC | CCCAG | CCCTA | CCCTT | CCCTC | CCCTG | CCCCA | CCCCT | CCCCC | CCCCG | CCCGA | CCCGT | CCCGC | CCCGG | | CCG | CCGAA | CCGAT | CCGAC | CCGAG | CCGTA | CCGTT | CCGTC | CCGTG | CCGCA | CCGCT | CCGCC | CCGCG | CCGGA | CCGGT | CCGGC | CCGGG | | CGA | CGAAA | CGAAT | CGAAC | CGAAG | CGATA | CGATT | CGATC | CGATG | CGACA | CGACT | CGACC | CGACG | CGAGA | CGAGT | CGAGC | CGAGG | | CGT | CGTAA | CGTAT | CGTAC | CGTAG | CGTTA | CGTTT | CGTTC | CGTTG | CGTCA | CGTCT | CGTCC | CGTCG | CGTGA | CGTGT | CGTGC | CGTGG | | CGC | CGCAA | CGCAT | CGCAC | CGCAG | CGCTA | CGCTT | CGCTC | CGCTG | CGCCA | CGCCT | CGCCC | CGCCG | CGCGA | CGCGT | CGCGC | CGCGG | | CGG | CGGAA | CGGAT | CGGAC | CGGAG | CGGTA | CGGTT | CGGTC | CGGTG | CGGCA | CGGCT | CGGCC | CGGCG | CGGGA | CGGGT | CGGGC | CGGGG | | GAA | GAAAA | GAAAT | GAAAC | GAAAG | GAATA | GAATT | GAATC | GAATG | GAACA | GAACT | GAACC | GAACG | GAAGA | GAAGT | GAAGC | GAAGG | | GAT | GATAA | GATAT | GATAC | GATAG | GATTA | GATTT | GATTC | GATTG | GATCA | GATCT | GATCC | GATCG | GATGA | GATGT | GATGC | GATGG | | GAC | GACAA | GACAT | GACAC | GACAG | GACTA | GACTT | GACTC | GACTG | GACCA | GACCT | GACCC | GACCG | GACGA | GACGT | GACGC | GACGG | | GAG | GAGAA | GAGAT | GAGAC | GAGAG | GAGTA | GAGTT | GAGTC | GAGTG | GAGCA | GAGCT | GAGCC | GAGCG | GAGGA | GAGGT | GAGGC | GAGGG | | GTA | GTAAA | GTAAT | GTAAC | GTAAG | GTATA | GTATT | GTATC | GTATG | GTACA | GTACT | GTACC | GTACG | GTAGA | GTAGT | GTAGC | GTAGG | | GTT | GTTAA | GTTAT | GTTAC | GTTAG | GTTTA | GTTTT | GTTTC | GTTTG | GTTCA | GTTCT | GTTCC | GTTCG | GTTGA | GTTGT | GTTGC | GTTGG | | GTC | GTCAA | GTCAT | GTCAC | GTCAG | GTCTA | GTCTT | GTCTC | GTCTG | GTCCA | GTCCT | GTCCC | GTCCG | GTCGA | GTCGT | GTCGC | GTCGG | | GTG | GTGAA | GTGAT | GTGAC | GTGAG | GTGTA | GTGTT | GTGTC | GTGTG | GTGCA | GTGCT | GTGCC | GTGCG | GTGGA | GTGGT | GTGGC | GTGGG | | GCA | GCAAA | GCAAT | GCAAC | GCAAG | GCATA | GCATT | GCATC | GCATG | GCACA | GCACT | GCACC | GCACG | GCAGA | GCAGT | GCAGC | GCAGG | | GCT | GCTAA | GCTAT | GCTAC | GCTAG | GCTTA | GCTTT | GCTTC | GCTTG | GCTCA | GCTCT | GCTCC | GCTCG | GCTGA | GCTGT | GCTGC | GCTGG | | GCC | GCCAA | GCCAT | GCCAC | GCCAG | GCCTA | GCCTT | GCCTC | GCCTG | GCCCA | GCCCT | GCCCC | GCCCG | GCCGA | GCCGT | GCCGC | GCCGG | | GCG | GCGAA | GCGAT | GCGAC | GCGAG | GCGTA | GCGTT | GCGTC | GCGTG | GCGCA | GCGCT | GCGCC | GCGCG | GCGGA | GCGGT | GCGGC | GCGGG | | GGA | GGAAA | GGAAT | GGAAC | GGAAG | GGATA | GGATT | GGATC | GGATG | GGACA | GGACT | GGACC | GGACG | GGAGA | GGAGT | GGAGC | GGAGG | | GGT | GGTAA | GGTAT | GGTAC | GGTAG | GGTTA | GGTTT | GGTTC | GGTTG | GGTCA | GGTCT | GGTCC | GGTCG | GGTGA | GGTGT | GGTGC | GGTGG | | GGC | GGCAA | GGCAT | GGCAC | GGCAG | GGCTA | GGCTT | GGCTC | GGCTG | GGCCA | GGCCT | GGCCC | GGCCG | GGCGA | GGCGT | GGCGC | GGCGG | | GGG | GGGAA | GGGAT | GGGAC | GGGAG | GGGTA | GGGTT | GGGTC | GGGTG | GGGCA | GGGCT | GGGCC | GGGCG | GGGGA | GGGGT | GGGGC | GGGGG | After filtering After filtering: read1: quality Value of each position will be shown on mouse over. After filtering: read1: base contents Value of each position will be shown on mouse over. After filtering: read1: KMER counting Darker background means larger counts. The count will be shown on mouse over. | | | | | | | | | | | | | | | | | | | --- | --- | --- | --- | --- | --- | --- | --- | --- | --- | --- | --- | --- | --- | --- | --- | --- | | | AA | AT | AC | AG | TA | TT | TC | TG | CA | CT | CC | CG | GA | GT | GC | GG | | AAA | AAAAA | AAAAT | AAAAC | AAAAG | AAATA | AAATT | AAATC | AAATG | AAACA | AAACT | AAACC | AAACG | AAAGA | AAAGT | AAAGC | AAAGG | | AAT | AATAA | AATAT | AATAC | AATAG | AATTA | AATTT | AATTC | AATTG | AATCA | AATCT | AATCC | AATCG | AATGA | AATGT | AATGC | AATGG | | AAC | AACAA | AACAT | AACAC | AACAG | AACTA | AACTT | AACTC | AACTG | AACCA | AACCT | AACCC | AACCG | AACGA | AACGT | AACGC | AACGG | | AAG | AAGAA | AAGAT | AAGAC | AAGAG | AAGTA | AAGTT | AAGTC | AAGTG | AAGCA | AAGCT | AAGCC | AAGCG | AAGGA | AAGGT | AAGGC | AAGGG | | ATA | ATAAA | ATAAT | ATAAC | ATAAG | ATATA | ATATT | ATATC | ATATG | ATACA | ATACT | ATACC | ATACG | ATAGA | ATAGT | ATAGC | ATAGG | | ATT | ATTAA | ATTAT | ATTAC | ATTAG | ATTTA | ATTTT | ATTTC | ATTTG | ATTCA | ATTCT | ATTCC | ATTCG | ATTGA | ATTGT | ATTGC | ATTGG | | ATC | ATCAA | ATCAT | ATCAC | ATCAG | ATCTA | ATCTT | ATCTC | ATCTG | ATCCA | ATCCT | ATCCC | ATCCG | ATCGA | ATCGT | ATCGC | ATCGG | | ATG | ATGAA | ATGAT | ATGAC | ATGAG | ATGTA | ATGTT | ATGTC | ATGTG | ATGCA | ATGCT | ATGCC | ATGCG | ATGGA | ATGGT | ATGGC | ATGGG | | ACA | ACAAA | ACAAT | ACAAC | ACAAG | ACATA | ACATT | ACATC | ACATG | ACACA | ACACT | ACACC | ACACG | ACAGA | ACAGT | ACAGC | ACAGG | | ACT | ACTAA | ACTAT | ACTAC | ACTAG | ACTTA | ACTTT | ACTTC | ACTTG | ACTCA | ACTCT | ACTCC | ACTCG | ACTGA | ACTGT | ACTGC | ACTGG | | ACC | ACCAA | ACCAT | ACCAC | ACCAG | ACCTA | ACCTT | ACCTC | ACCTG | ACCCA | ACCCT | ACCCC | ACCCG | ACCGA | ACCGT | ACCGC | ACCGG | | ACG | ACGAA | ACGAT | ACGAC | ACGAG | ACGTA | ACGTT | ACGTC | ACGTG | ACGCA | ACGCT | ACGCC | ACGCG | ACGGA | ACGGT | ACGGC | ACGGG | | AGA | AGAAA | AGAAT | AGAAC | AGAAG | AGATA | AGATT | AGATC | AGATG | AGACA | AGACT | AGACC | AGACG | AGAGA | AGAGT | AGAGC | AGAGG | | AGT | AGTAA | AGTAT | AGTAC | AGTAG | AGTTA | AGTTT | AGTTC | AGTTG | AGTCA | AGTCT | AGTCC | AGTCG | AGTGA | AGTGT | AGTGC | AGTGG | | AGC | AGCAA | AGCAT | AGCAC | AGCAG | AGCTA | AGCTT | AGCTC | AGCTG | AGCCA | AGCCT | AGCCC | AGCCG | AGCGA | AGCGT | AGCGC | AGCGG | | AGG | AGGAA | AGGAT | AGGAC | AGGAG | AGGTA | AGGTT | AGGTC | AGGTG | AGGCA | AGGCT | AGGCC | AGGCG | AGGGA | AGGGT | AGGGC | AGGGG | | TAA | TAAAA | TAAAT | TAAAC | TAAAG | TAATA | TAATT | TAATC | TAATG | TAACA | TAACT | TAACC | TAACG | TAAGA | TAAGT | TAAGC | TAAGG | | TAT | TATAA | TATAT | TATAC | TATAG | TATTA | TATTT | TATTC | TATTG | TATCA | TATCT | TATCC | TATCG | TATGA | TATGT | TATGC | TATGG | | TAC | TACAA | TACAT | TACAC | TACAG | TACTA | TACTT | TACTC | TACTG | TACCA | TACCT | TACCC | TACCG | TACGA | TACGT | TACGC | TACGG | | TAG | TAGAA | TAGAT | TAGAC | TAGAG | TAGTA | TAGTT | TAGTC | TAGTG | TAGCA | TAGCT | TAGCC | TAGCG | TAGGA | TAGGT | TAGGC | TAGGG | | TTA | TTAAA | TTAAT | TTAAC | TTAAG | TTATA | TTATT | TTATC | TTATG | TTACA | TTACT | TTACC | TTACG | TTAGA | TTAGT | TTAGC | TTAGG | | TTT | TTTAA | TTTAT | TTTAC | TTTAG | TTTTA | TTTTT | TTTTC | TTTTG | TTTCA | TTTCT | TTTCC | TTTCG | TTTGA | TTTGT | TTTGC | TTTGG | | TTC | TTCAA | TTCAT | TTCAC | TTCAG | TTCTA | TTCTT | TTCTC | TTCTG | TTCCA | TTCCT | TTCCC | TTCCG | TTCGA | TTCGT | TTCGC | TTCGG | | TTG | TTGAA | TTGAT | TTGAC | TTGAG | TTGTA | TTGTT | TTGTC | TTGTG | TTGCA | TTGCT | TTGCC | TTGCG | TTGGA | TTGGT | TTGGC | TTGGG | | TCA | TCAAA | TCAAT | TCAAC | TCAAG | TCATA | TCATT | TCATC | TCATG | TCACA | TCACT | TCACC | TCACG | TCAGA | TCAGT | TCAGC | TCAGG | | TCT | TCTAA | TCTAT | TCTAC | TCTAG | TCTTA | TCTTT | TCTTC | TCTTG | TCTCA | TCTCT | TCTCC | TCTCG | TCTGA | TCTGT | TCTGC | TCTGG | | TCC | TCCAA | TCCAT | TCCAC | TCCAG | TCCTA | TCCTT | TCCTC | TCCTG | TCCCA | TCCCT | TCCCC | TCCCG | TCCGA | TCCGT | TCCGC | TCCGG | | TCG | TCGAA | TCGAT | TCGAC | TCGAG | TCGTA | TCGTT | TCGTC | TCGTG | TCGCA | TCGCT | TCGCC | TCGCG | TCGGA | TCGGT | TCGGC | TCGGG | | TGA | TGAAA | TGAAT | TGAAC | TGAAG | TGATA | TGATT | TGATC | TGATG | TGACA | TGACT | TGACC | TGACG | TGAGA | TGAGT | TGAGC | TGAGG | | TGT | TGTAA | TGTAT | TGTAC | TGTAG | TGTTA | TGTTT | TGTTC | TGTTG | TGTCA | TGTCT | TGTCC | TGTCG | TGTGA | TGTGT | TGTGC | TGTGG | | TGC | TGCAA | TGCAT | TGCAC | TGCAG | TGCTA | TGCTT | TGCTC | TGCTG | TGCCA | TGCCT | TGCCC | TGCCG | TGCGA | TGCGT | TGCGC | TGCGG | | TGG | TGGAA | TGGAT | TGGAC | TGGAG | TGGTA | TGGTT | TGGTC | TGGTG | TGGCA | TGGCT | TGGCC | TGGCG | TGGGA | TGGGT | TGGGC | TGGGG | | CAA | CAAAA | CAAAT | CAAAC | CAAAG | CAATA | CAATT | CAATC | CAATG | CAACA | CAACT | CAACC | CAACG | CAAGA | CAAGT | CAAGC | CAAGG | | CAT | CATAA | CATAT | CATAC | CATAG | CATTA | CATTT | CATTC | CATTG | CATCA | CATCT | CATCC | CATCG | CATGA | CATGT | CATGC | CATGG | | CAC | CACAA | CACAT | CACAC | CACAG | CACTA | CACTT | CACTC | CACTG | CACCA | CACCT | CACCC | CACCG | CACGA | CACGT | CACGC | CACGG | | CAG | CAGAA | CAGAT | CAGAC | CAGAG | CAGTA | CAGTT | CAGTC | CAGTG | CAGCA | CAGCT | CAGCC | CAGCG | CAGGA | CAGGT | CAGGC | CAGGG | | CTA | CTAAA | CTAAT | CTAAC | CTAAG | CTATA | CTATT | CTATC | CTATG | CTACA | CTACT | CTACC | CTACG | CTAGA | CTAGT | CTAGC | CTAGG | | CTT | CTTAA | CTTAT | CTTAC | CTTAG | CTTTA | CTTTT | CTTTC | CTTTG | CTTCA | CTTCT | CTTCC | CTTCG | CTTGA | CTTGT | CTTGC | CTTGG | | CTC | CTCAA | CTCAT | CTCAC | CTCAG | CTCTA | CTCTT | CTCTC | CTCTG | CTCCA | CTCCT | CTCCC | CTCCG | CTCGA | CTCGT | CTCGC | CTCGG | | CTG | CTGAA | CTGAT | CTGAC | CTGAG | CTGTA | CTGTT | CTGTC | CTGTG | CTGCA | CTGCT | CTGCC | CTGCG | CTGGA | CTGGT | CTGGC | CTGGG | | CCA | CCAAA | CCAAT | CCAAC | CCAAG | CCATA | CCATT | CCATC | CCATG | CCACA | CCACT | CCACC | CCACG | CCAGA | CCAGT | CCAGC | CCAGG | | CCT | CCTAA | CCTAT | CCTAC | CCTAG | CCTTA | CCTTT | CCTTC | CCTTG | CCTCA | CCTCT | CCTCC | CCTCG | CCTGA | CCTGT | CCTGC | CCTGG | | CCC | CCCAA | CCCAT | CCCAC | CCCAG | CCCTA | CCCTT | CCCTC | CCCTG | CCCCA | CCCCT | CCCCC | CCCCG | CCCGA | CCCGT | CCCGC | CCCGG | | CCG | CCGAA | CCGAT | CCGAC | CCGAG | CCGTA | CCGTT | CCGTC | CCGTG | CCGCA | CCGCT | CCGCC | CCGCG | CCGGA | CCGGT | CCGGC | CCGGG | | CGA | CGAAA | CGAAT | CGAAC | CGAAG | CGATA | CGATT | CGATC | CGATG | CGACA | CGACT | CGACC | CGACG | CGAGA | CGAGT | CGAGC | CGAGG | | CGT | CGTAA | CGTAT | CGTAC | CGTAG | CGTTA | CGTTT | CGTTC | CGTTG | CGTCA | CGTCT | CGTCC | CGTCG | CGTGA | CGTGT | CGTGC | CGTGG | | CGC | CGCAA | CGCAT | CGCAC | CGCAG | CGCTA | CGCTT | CGCTC | CGCTG | CGCCA | CGCCT | CGCCC | CGCCG | CGCGA | CGCGT | CGCGC | CGCGG | | CGG | CGGAA | CGGAT | CGGAC | CGGAG | CGGTA | CGGTT | CGGTC | CGGTG | CGGCA | CGGCT | CGGCC | CGGCG | CGGGA | CGGGT | CGGGC | CGGGG | | GAA | GAAAA | GAAAT | GAAAC | GAAAG | GAATA | GAATT | GAATC | GAATG | GAACA | GAACT | GAACC | GAACG | GAAGA | GAAGT | GAAGC | GAAGG | | GAT | GATAA | GATAT | GATAC | GATAG | GATTA | GATTT | GATTC | GATTG | GATCA | GATCT | GATCC | GATCG | GATGA | GATGT | GATGC | GATGG | | GAC | GACAA | GACAT | GACAC | GACAG | GACTA | GACTT | GACTC | GACTG | GACCA | GACCT | GACCC | GACCG | GACGA | GACGT | GACGC | GACGG | | GAG | GAGAA | GAGAT | GAGAC | GAGAG | GAGTA | GAGTT | GAGTC | GAGTG | GAGCA | GAGCT | GAGCC | GAGCG | GAGGA | GAGGT | GAGGC | GAGGG | | GTA | GTAAA | GTAAT | GTAAC | GTAAG | GTATA | GTATT | GTATC | GTATG | GTACA | GTACT | GTACC | GTACG | GTAGA | GTAGT | GTAGC | GTAGG | | GTT | GTTAA | GTTAT | GTTAC | GTTAG | GTTTA | GTTTT | GTTTC | GTTTG | GTTCA | GTTCT | GTTCC | GTTCG | GTTGA | GTTGT | GTTGC | GTTGG | | GTC | GTCAA | GTCAT | GTCAC | GTCAG | GTCTA | GTCTT | GTCTC | GTCTG | GTCCA | GTCCT | GTCCC | GTCCG | GTCGA | GTCGT | GTCGC | GTCGG | | GTG | GTGAA | GTGAT | GTGAC | GTGAG | GTGTA | GTGTT | GTGTC | GTGTG | GTGCA | GTGCT | GTGCC | GTGCG | GTGGA | GTGGT | GTGGC | GTGGG | | GCA | GCAAA | GCAAT | GCAAC | GCAAG | GCATA | GCATT | GCATC | GCATG | GCACA | GCACT | GCACC | GCACG | GCAGA | GCAGT | GCAGC | GCAGG | | GCT | GCTAA | GCTAT | GCTAC | GCTAG | GCTTA | GCTTT | GCTTC | GCTTG | GCTCA | GCTCT | GCTCC | GCTCG | GCTGA | GCTGT | GCTGC | GCTGG | | GCC | GCCAA | GCCAT | GCCAC | GCCAG | GCCTA | GCCTT | GCCTC | GCCTG | GCCCA | GCCCT | GCCCC | GCCCG | GCCGA | GCCGT | GCCGC | GCCGG | | GCG | GCGAA | GCGAT | GCGAC | GCGAG | GCGTA | GCGTT | GCGTC | GCGTG | GCGCA | GCGCT | GCGCC | GCGCG | GCGGA | GCGGT | GCGGC | GCGGG | | GGA | GGAAA | GGAAT | GGAAC | GGAAG | GGATA | GGATT | GGATC | GGATG | GGACA | GGACT | GGACC | GGACG | GGAGA | GGAGT | GGAGC | GGAGG | | GGT | GGTAA | GGTAT | GGTAC | GGTAG | GGTTA | GGTTT | GGTTC | GGTTG | GGTCA | GGTCT | GGTCC | GGTCG | GGTGA | GGTGT | GGTGC | GGTGG | | GGC | GGCAA | GGCAT | GGCAC | GGCAG | GGCTA | GGCTT | GGCTC | GGCTG | GGCCA | GGCCT | GGCCC | GGCCG | GGCGA | GGCGT | GGCGC | GGCGG | | GGG | GGGAA | GGGAT | GGGAC | GGGAG | GGGTA | GGGTT | GGGTC | GGGTG | GGGCA | GGGCT | GGGCC | GGGCG | GGGGA | GGGGT | GGGGC | GGGGG |

fastp -i ../sources/combined\_short\_reads.fastq.gz -w 16

fastp 0.23.4, at 2023-08-01 17:46:56
